# Human microcephaly ASPM protein is a spindle pole-focusing factor that functions redundantly with CDK5RAP2

**DOI:** 10.1101/102236

**Authors:** Elsa A. Tungadi, Ami Ito, Tomomi Kiyomitsu, Gohta Goshima

## Abstract

Nonsense mutations in the *ASPM* gene have been most frequently identified among familial microcephaly patients. Depletion of the *Drosophila* orthologue causes spindle pole unfocusing during mitosis in multiple cell types. However, it remains unknown whether human ASPM has a similar function. Here, using CRISPR-based gene knockout (KO) and RNA interference combined with auxin-inducible degron, we show that ASPM functions in spindle pole organisation during mitotic metaphase redundantly with another microcephaly protein CDK5RAP2 (also called CEP215) in human tissue culture cells. Deletion of the *ASPM* gene alone did not affect spindle morphology or mitotic progression. However, when the pericentriolar material protein CDK5RAP2 was depleted in *ASPM* KO cells, spindle poles were unfocused during prometaphase and anaphase onset was significantly delayed. The phenotypic analysis of CDK5RAP2-depleted cells suggested that the pole-focusing function of CDK5RAP2 is independent of its known function to localise the kinesin-14 motor HSET or activate the *γ*-tubulin complex. Finally, a hypomorphic mutation identified in ASPM microcephaly patients similarly caused spindle pole unfocusing in the absence of CDK5RAP2, suggesting a possible link between spindle pole disorganisation and microcephaly.

## Introduction

The most common cause of autosomal recessive primary microcephaly is a homozygous mutation of the *abnormal spindle-like microcephaly-associated* (*ASPM*) gene (Bond et al., 2002, Tan et al., 2014, Bond et al., 2003, Abdel-Hamid et al., 2016). *ASPM* was originally identified in *Drosophila*, as the orthologue *Asp* (abnormal spindle), whose mutation results in abnormal spindle formation (Ripoll et al., 1985). In the absence of Asp, centrosomes are detached from the main body of the spindle, and spindle microtubules (MTs) are unfocused at the pole (Saunders et al., 1997, Wakefield et al., 2001, Morales-Mulia and Scholey, 2005, Ito and Goshima, 2015, Schoborg et al., 2015). Asp is concentrated at the spindle pole, the area enriched with the spindle MT minus ends. The current model is that Asp binds directly to the spindle MT ends using the middle region containing calponin homology domains and cross-links them each other using the C-terminal domain, thus postulating Asp as a critical pole-focusing factor (Ito and Goshima, 2015). Recent studies have also shown that *asp* mutant flies have reduced brain size (Rujano et al., 2013, Schoborg et al., 2015), which was at least partly attributed to chromosome mis-segregation associated with unfocused spindle poles (Rujano et al., 2013).

The cellular function of human ASPM has been evaluated using RNA interference (RNAi)-mediated knockdown. Small interfering RNA (siRNA)-based knockdown in U2OS cells led to spindle misorientation, cytokinesis failure, reduction in mitotic cells, and apoptosis (Higgins et al., 2010), possibly through interactions with the citron kinase (Gai et al., 2016, Paramasivam et al., 2007). Another study observed downregulation of BRCA1 protein upon ASPM knockdown (Zhong et al., 2005). To the best of our knowledge, the pole-focusing defect commonly observed upon Asp depletion in *Drosophila* has not been reported in studies of human ASPM knockdown. However, RNAi has a general limitation in that residual protein expression might suffice to fulfil the function of the target protein. Moreover, none of the previous RNAi studies of ASPM involved a rescue experiment in which full-length ASPM was ectopically expressed after endogenous ASPM depletion, leaving the possibility that some of the observed phenotypes were derived from off-target effects of the siRNAs utilised. The effect of mutations identified in microcephaly patients has also not been assessed using a cell culture model. Nevertheless, the lack of the spindle pole phenotype is surprising, given the result obtained in *Drosophila*.

In this study, we used CRISPR/Cas9-based knockout (KO) as well as RNAi to decipher the mitotic function of ASPM in the human HCT116 cell line. Our data provide the first demonstration that human ASPM is an important factor in spindle pole organisation, working redundantly with the centrosomal protein CDK5RAP2. Furthermore, a mutation identified in microcephaly patients similarly impaired this function of ASPM.

## Results

### No mitotic abnormality upon *ASPM* KO in the human HCT116 cell line

To precisely assess the spindle pole-organising function of human ASPM, we utilised the human colorectal cancer cell line HCT116, which has a stable diploid karyotype, assembles well-focused bipolar spindles, and is highly amenable to CRISPR/Cas9-based genome editing and other gene perturbation techniques such as RNAi and auxin-inducible degron (AID) (Natsume et al., 2016). We aimed to delete the *ASPM* gene in HCT116 cells using CRISPR/Cas9-based genome editing. Since it was reported that all ASPM isoforms are translated from a common start codon in exon 1 (Kouprina et al., 2005), we aimed to select homozygous KO lines by inserting a drug-resistant marker in exon 1 and deleting the start codon (Fig. S2A, B). Immunofluorescence microscopy using anti-ASPM antibody indicated that ASPM was significantly depleted in the selected lines (termed KO^Exon1^), confirming successful editing (Fig. S2C). However, in the course of this project, we noticed that some residual protein expression was still detected with this antibody when the image contrast was further enhanced (Fig. S2C, quantification is presented in Fig. 4B). Therefore, we also selected the cell lines in which the entire open reading frame was replaced by a marker gene cassette (Fig. 1A, S1). The polar staining completely disappeared in the complete KO line (termed KO^C^; Fig. 1B, 4B). In either KO line, time-lapse microscopy of MTs did not reveal any abnormality in mitotic progression and spindle formation, suggesting that *ASPM* is not an essential gene for mitosis in this cell line (Fig. 1C, D; S2D, E; and Movie 1).

**Figure 1.**
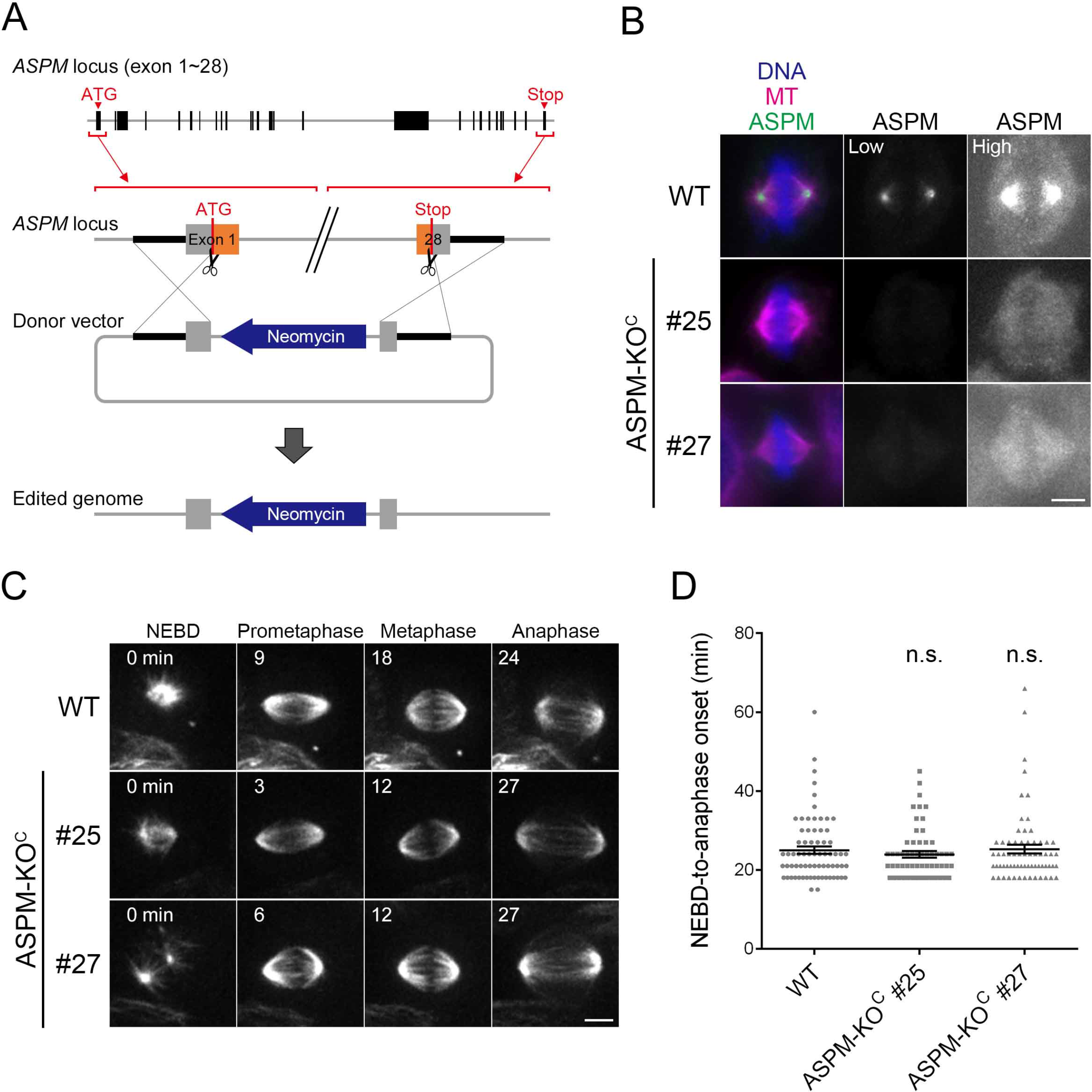
The *ASPM* KO line shows no defects in the spindle pole. **(A, B)** Construction of the complete *ASPM* KO line (KO^c^), in which the entire *ASPM* open reading frame was replaced by the neomycin-resistant marker. **(B)** Complete *ASPM* KO (KO^c^) lines did not give rise to pole staining with anti-ASPM antibody. Images with highly enhanced signals are presented on the right. **(C)** Spindle dynamics of *ASPM* KO^c^lines. MTs were visualized by SiR-tubulin staining. Images were acquired with 5 z-sections (separated by 3 µm) and displayed after maximum projection. No abnormality was detected. Images of two clones (#25 and #27) are shown. See also Movie **(D)** Mitotic duration with or without ASPM. Data are mean± SEM. WT; 25 ± 1 (n = 71), KO #25; 24 ± 1 (n = 59, p > 0.4), KO #27; 25 ± 1 (n = 66, p > 0.8). n.s. stands for ‘not significant’. Bars, 5 µm.

### Spindle poles were unfocused upon double depletion of ASPM and CDK5RAP2

The centrosome constitutes the focal point of spindle MTs at the pole in animal somatic cells. However, in flies and mammals, acentrosomal spindles with two focused poles can be assembled after centrosomal protein depletion/inhibition, indicating that centrosomes are not a prerequisite for pole focusing (Wong et al., 2015, Megraw et al., 2001). Nevertheless, centrosomes play a supportive role in spindle MT focusing, perhaps by supplying additional MTs for MT cross-linking motors and MAPs for the interaction (Goshima et al., 2005, Ito and Goshima, 2015, Silk et al., 2009, Wakefield et al., 2001, Mountain et al., 1999, Heald et al., 1997, Baumbach et al., 2015, Chavali et al., 2016). We hypothesised that lack of the pole unfocusing phenotype in the absence of ASPM may also be partly due to the presence of strong centrosomes in this cell line.

To test this possibility, we aimed to deplete a core centrosomal protein. In *Drosophila*, the pericentriolar material protein Centrosomin (Cnn) is critical for the centrosomal enrichment of the γ-tubulin ring complex (γ-TuRC), and thereby centrosomal MT formation (Megraw et al., 2001). In mammals, the Cnn orthologue CDK5RAP2 (also known as CEP215) is a microcephaly protein that activates γ-TuRC *in vitro* (Choi et al., 2010, Bond et al., 2005). In U2OS cells, astral MTs from centrosomes were dramatically reduced in the absence of CDK5RAP2, consistent with its biochemical activity and the *Drosophila* phenotype (Fong et al., 2008). However, in other human cell types (B lymphocytes, BT-546 breast cancer cells, HeLa cells), such phenotypes have not been observed; instead, centrosome detachment was reported. In these cells, delocalisation of the kinesin-14 motor HSET from the centrosome was attributed to this phenotype (Chavali et al., 2016).

In the current study, we first depleted CDK5RAP2 by RNAi in the complete ASPM KO^C^ line (Fig. 2A– C, Movie 2). In cells with intact ASPM, bipolar spindles with focused poles were observed in >90% of cases, regardless of the presence or absence of endogenous CDK5RAP2 (tagged with mCherry), and most cells proceeded into anaphase with only a slight delay, even when CDK5RAP2-mCherry signals were hardly detected (1.2 ± 0.0-fold increase, NEBD to anaphase onset, n = 3 experiments). In sharp contrast, when CDK5RAP2 was depleted in the *ASPM* KO^C^ background, >60% of the cells exhibited unfocused spindle poles; spindle MTs were not persistently associated with the pole during metaphase (Fig. 2A, B). In these cells, anaphase onset was drastically delayed (4.3 ± 1.5-fold increase in prometaphase/metaphase duration, n = 3 experiments; Fig. 2C).

**Figure 2.**
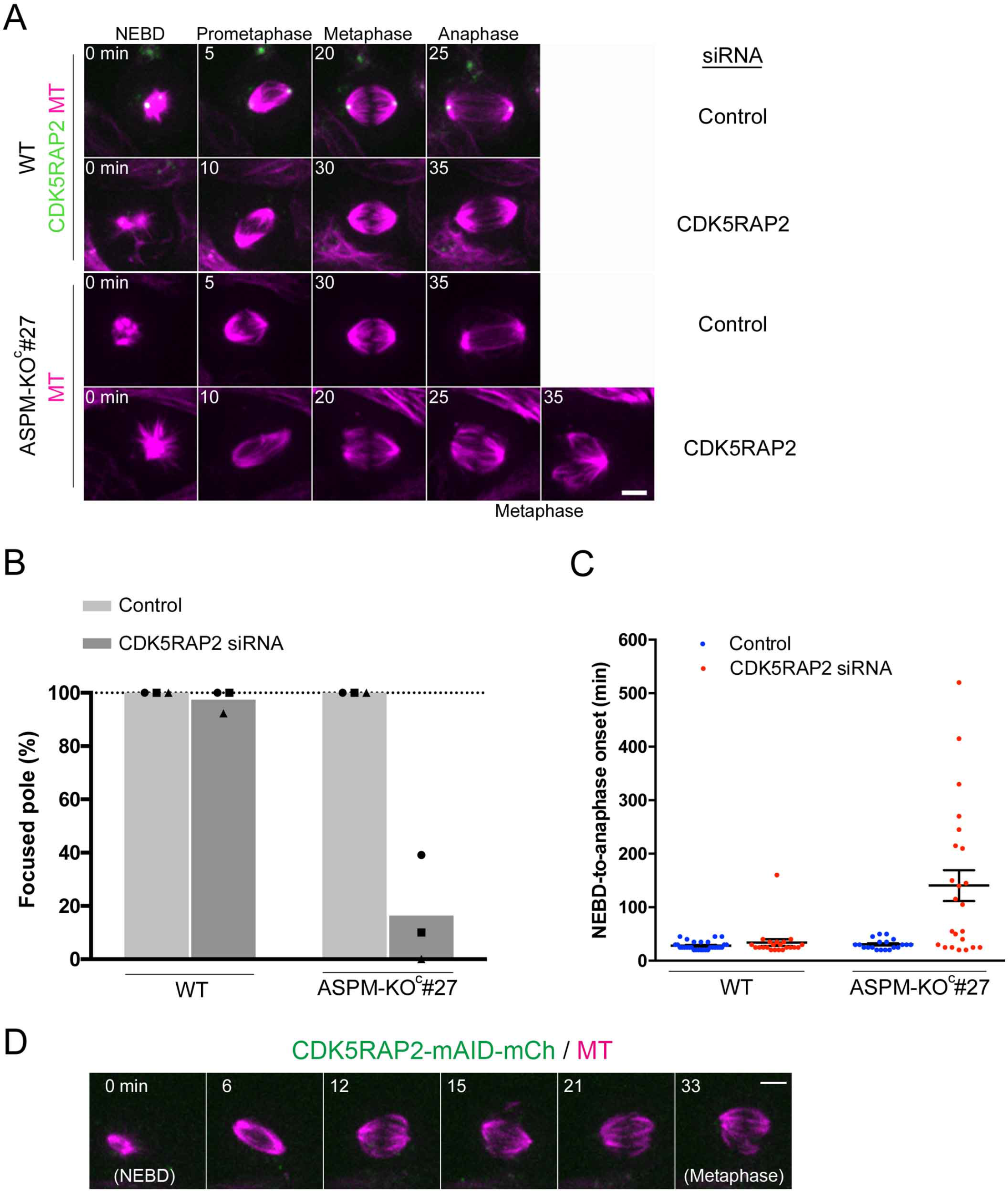
Spindle pole unfocusing and mitotic delay following CDK5RAP2 depletion in the *ASPM* KO line. **(A, B)** CDK5RAP2 was depleted by RNAi in the ASPM KOc #27 line and the control CDK5RAP2-mAID-mCherry line. Images were acquired with 5 z-sections (separated by 3 µm) and displayed after maximum projection. The values of each experiment are indicated by dots, whereas the mean values of three independent experiments are indicated by bars. The number of spindles analysed were (10, 30, 10], (13, 22, 10], (10, 20, 10], and [10, 23, 13] (from left to right). Unfocused poles were dramatically increased in the absence of CDK5RAP2 and ASPM (p < 0.02 vs. luciferase RNAi control). **(C)** Mitotic duration with and without CDK5RAP2 (mean ± SEM). CDK5RAP2 RNAi treatment caused a dramatic increase in mitotic duration in ASPM-KOc#27 cells compared to untreated KO cells (p < 0.0001; n = 30, 22, 20, 23 from left to right). Bar, 5 µm. **(D)** Spindle pole unfocusing in the ASPM KOExon ^1^ line following CDK5RAP2 depletion. Treatment with Dox and IAA induced the degradation of CDK5RAP2-mAID-mCherry. Images were acquired with 5 z-sections (separated by 3 µm) and displayed after maximum projection. Two independent KOExon ^1^ lines (#3 and #20) were analysed three times. The NEBD-to-anaphase duration was dramatically increased in these cells (161 ± 20 min [±SEM, n = 61]). Bars, 5 µm.

As another means to deplete CDK5RAP2, we introduced the auxin-inducible degron (AID) system, which allows the acute degradation of a target protein (Nishimura et al., 2009). We tagged endogenous CDK5RAP2 with mCherry and mini-AID (mAID) sequences in cells possessing the gene encoding *Oryza sativa* (Os) TIR1, the auxin responsive F-box protein that constitutes a functional SCF ubiquitin ligase in human cells (Fig. S3A–C). In this cell line, OsTIR1 protein was induced by doxycycline hyclate (Dox) (Natsume et al., 2016), and CDK5RAP2-mAID-mCherry was subsequently degraded upon auxin (indole-3-acetic acid; IAA) addition to the culture medium (Fig. S3D). We deleted the first exon of *ASPM* in this line and observed spindle dynamics after 24 h of auxin treatment. We observed an identical phenotype to that observed upon RNAi (Fig. 2D; unfocusing was observed in 82% (n = 68) or 86% (n = 72) of KO^Exon1^ #3 and KO^Exon1^ #20 lines, respectively). We concluded that ASPM is required for pole focusing when the CDK5RAP2 function is compromised.

To investigate if the synthetic polar phenotype is attributed to γ-tubulin delocalisation and reduction of astral MTs (Fong et al., 2008) or HSET delocalisation (Chavali et al., 2016) in the absence of CDK5RAP2, we immunostained for γ-tubulin, HSET, and MTs after CDK5RAP2 depletion (Fig. S4). Contrary to the result in U2OS cells, we clearly observed γ-tubulin at the centrosome with an apparently normal number of associated MTs (Fig. S4A). Furthermore, HSET was uniformly localised at the spindle, which was indistinguishable between control and CDK5RAP2-depleted cells (Fig. S4B). We also assessed ASPM localisation after CDK5RAP2 depletion, and found that ASPM was similarly enriched at the pole in the absence of CDK5RAP2 (Fig. S4C).

### No synthetic pole phenotypes between HSET and ASPM

A previous study on *Drosophila* S2 cells showed that spindle MT focusing at the pole was ensured by the independent actions of kinesin-14 and Asp (Ito and Goshima, 2015). We treated control and *ASPM* KO^C^ cells with siRNA against HSET. Immunofluorescence microscopy indicated that HSET was efficiently knocked down at 48 h, as was expected from a previous study (Cai et al., 2009) (Fig. 3A). However, in time-lapse observation of mitosis, unfocused poles were observed at a similarly low frequency between control (9%, n = 34) and HSET-depleted cells (13%, n = 31), and no mitotic delay was observed without HSET (27 ± 2 min [SEM], n = 11; Fig. 3C). These results indicated that pole focusing was achieved in the absence of ASPM and HSET in this cell line.

**Figure 3.**
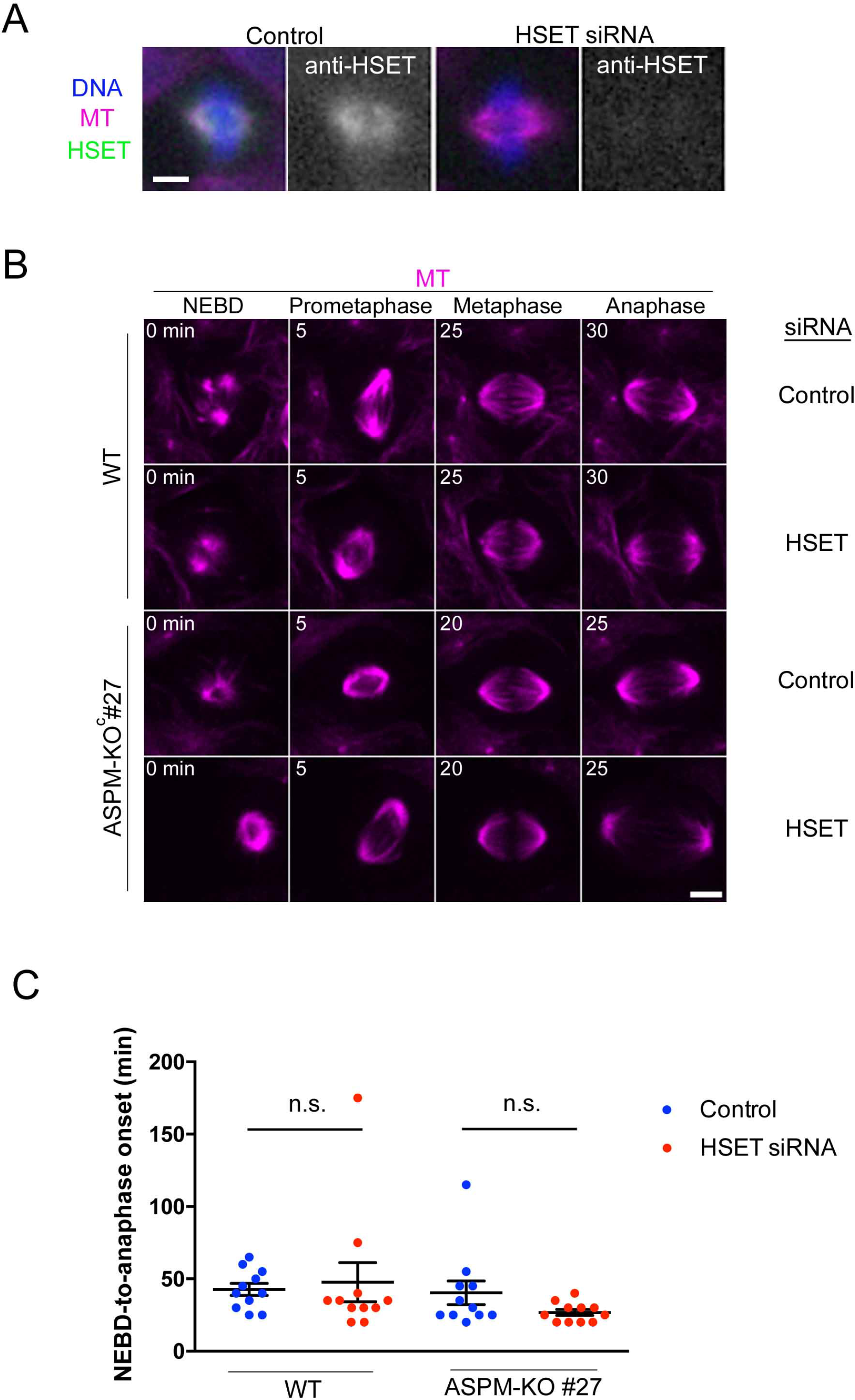
HSET inhibition does not provoke synthetic pole disorganisation with ASPM. **(A)** RNAi depletion of HSET was confirmed with immunofluorescence microscopy. **(B)** Spindle dynamics of ASPM KOc lines in the presence or absence of HSET. MTs were visualized by SiR-tubulin staining. Images were acquired with 5 z-sections (separated by 3 µm) and displayed after maximum projection. No abnormalities were detected in the double ASPM/HSET-depleted line. **(C)** Mitotic duration (mean ± SEM, n = 11 each). Bars, 5 µm.

### RNAi depletion of ASPM phenocopies CRISPR-based KO

One possible reason for the lack of mitotic defects in the *ASPM* KO line is that suppressor mutations were introduced into another locus during the few weeks of KO line selection. To exclude this possibility, we depleted ASPM more rapidly from the cell using RNAi. We prepared four independent siRNA constructs, one of which was previously reported to give rise to abnormal spindle orientation and M-phase reduction in U2OS cells after 72 h (Higgins et al., 2010) (Table S1). Immunostaining using anti-ASPM antibody indicated that all constructs efficiently knocked down ASPM protein after 48 h in HCT116 cells; the intensity of the residual ASPM signal was similar among the four constructs and to that of the KO^Exon1^ line (Fig. 4A, B). We observed a strong toxic effect for one of the constructs (siRNA #2) and mitotic acceleration for another (siRNA #3), suggesting that these constructs had off-target effects on important proteins other than ASPM (Table S1). However, cell proliferation, spindle morphogenesis, and mitotic progression were not defective for the remaining two constructs, reminiscent of the results observed in the KO line. These results strongly suggest that rapid ASPM depletion also has little impact on spindle organisation.

**Figure 4.**
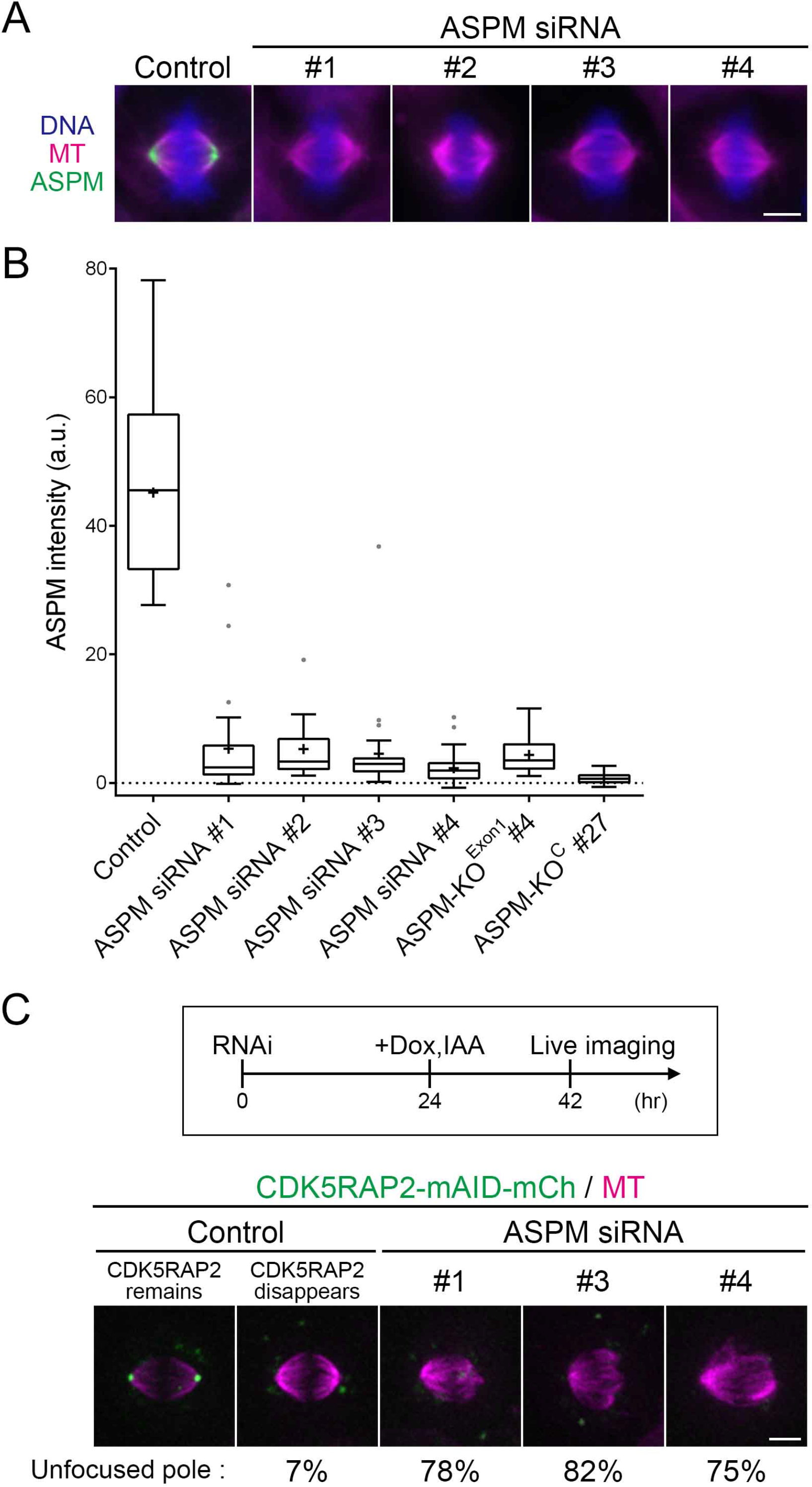
RNAi-mediated knockdown phenocopies KO. **(A, B)** Efficient knockdown of ASPM using four independent RNAi constructs as indicated by immunofluorescence microscopy (A) and the signal intensity measurement (B). The data are shown as a box-and-whisker plot, with the median (line in the middle of box), upper and lower quartiles (the ends of box), maximum and minimum (whiskers), outlier (dots), and mean (marked ‘+’) values presented; n = 28, 25, 13, 27, 32, 30, 28 (from left to right). Experiments were performed twice, and the result from one experiment is displayed in the graph. **(C)** Spindle pole unfocusing by combined ASPM knockdown and CDK5RAP2 degradation. Two control images(± CDK5RAP2-mCherry signals) are also displayed. Images were acquired with 5 z-sections (separated by 3 µm) and displayed after maximum projection; n = 39, 9, 11, 12 (left to right). Experiments were performed twice, and the result from one experiment is displayed. The result of the second experiment was as follows: 5% (n = 21), 100% (3), 64% (11), and 70% (16). Bars, 5 µm.

To test whether the synthetic phenotype with CDK5RAP2 depletion is also observed for *ASPM* RNAi, we treated the cells with three *ASPM* siRNAs (excluding the toxic siRNA construct) and then degraded CDK5RAP2 with the AID system. Time-lapse imaging indicated that pole unfocusing was frequently caused by CDK5RAP2 degradation for each siRNA construct (Fig. 4C). These results support the conclusion that ASPM becomes critical when the CDK5RAP2 function is compromised.

### Microcephaly mutation in *ASPM* causes pole unfocusing in the absence of CDK5RAP2

Homozygous mutations in the *ASPM* gene have been identified at various positions in microcephaly patients, almost all of which introduce a premature stop codon in the gene (Bond et al., 2002, Tan et al., 2014, Bond et al., 2003, Nicholas et al., 2009, Abdel-Hamid et al., 2016). The two most C-terminal mutations identified in three previous studies were located in front of the HEAT repeat motif: a nonsense mutation at amino acid (aa) 3,233 and a 1-bp deletion at aa 3,252, which introduces a stop codon at aa 3,261 (Fig. 5A). To test the effect of microcephaly mutations in spindle pole organisation, we inserted mClover next to the aa 3,232 residue to closely mimic a patient mutation (Fig. 5A, Fig. S5A, C); we selected this mutation, and not the most C-terminal mutation, as the former site was located next to the CRISPR-friendly sequences. As the control, mClover was inserted at the C-terminus of the *ASPM* gene (Fig. S5B, C). As expected, several replacement lines were obtained for both constructs. ASPM [1– 3,232]-mClover showed indistinguishable localisation at the pole from that of the full-length ASPM-mClover, consistent with the case of *Drosophila* Asp where the HEAT repeat was shown to be unnecessary for MT minus-end localisation (Ito and Goshima, 2015). However, when CDK5RAP2 was depleted by the AID system, we observed a severe pole unfocusing phenotype and mitotic delay for the ASPM [1–3,232]-mClover lines, reminiscent of the results for the *ASPM* KO line (Fig. 5B, C; Movie 3; mitotic duration, 298 ± 48 min, ± SEM, n = 11 for line#17 and 166 ± 49 min, n = 10 for line#19). Thus, the mutation that mimics that of microcephaly patients impaired the function of ASPM in pole organisation. Since other nonsense mutations and insertions/deletions in patients were identified upstream of the introduced mutation, a similar pole-unfocusing phenotype in the absence of CDK5RAP2 is expected to be observed for all cases.

**Figure 5.**
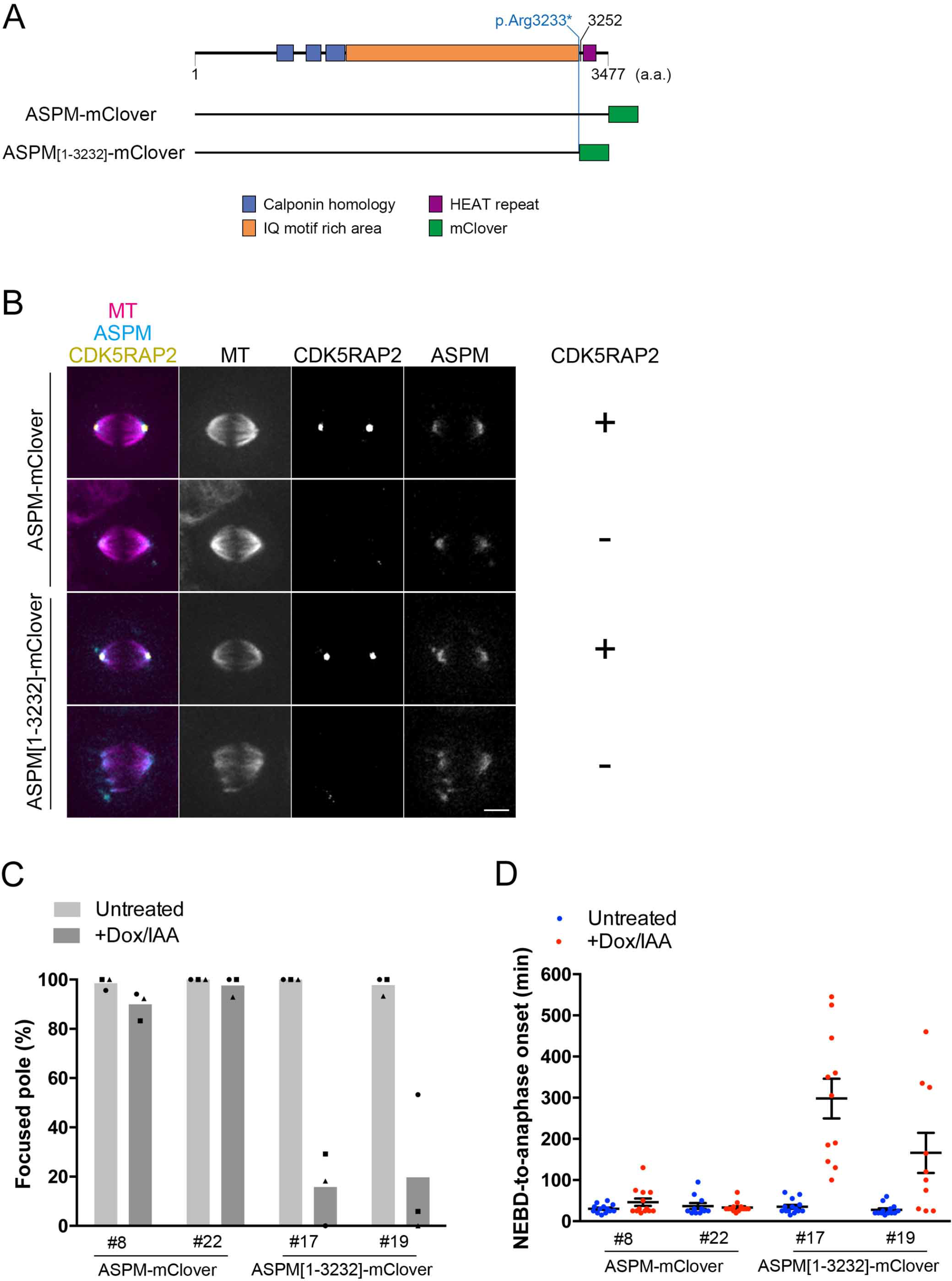
Hypomorphic microcephaly mutation impairs the spindle pole-focusing function of ASPM. **(A)** mClover insertion after the aa 3,232 residue mimics the second-most C-terminally truncated protein identified among the microcephaly patients (Tan et al., 2014, Bond et al., 2003, Abdel-Hamid et al., 2016). As a control, mClover was tagged to the C-terminusof the full-length (FL) ASPM protein. **(B, C)** Spindle pole unfocusing and mitotic delay caused by ASPM truncation in the absence of CDK5RAP2. Treatment with Dox and IAA induced the degradation of CDK5RAP2-mAID-mCherry . Two independent truncation lines (clones #17 and #19) and control full-length ASPM-mClover lines (clones #8 and #22) were analysed. See also Movie 3. Images were acquired with 5 z-sections (separated by 3 µm) and displayed after maximum projection. The values of each experiment are indicated by dots, whereas the mean values of three independent experiments are indicated by bars. The numbers of spindles analysed were [23, 8, 13], [17, 30, 13], [7, 26, 11], [10, 9, 14], [9, 6, 14], [14, 24, 11], [8, 15, 13], and [15, 17, 10] (from left to right). **(D)** Mitotic duration. Data are mean± SEM. ASPM [1-3232]#17, 298 ± 48 min, n = 11 (p < 0.0001 against untreated FL#8, 30 ± 3 min, n = 13) and ASPM [1-3232]#19, 166 ± 49 min, n = 10 (p < 0.0044). Bar, 5 µm.

## Discussion

### Spindle pole focusing by ASPM

The results described above established that human ASPM is a spindle pole-focusing factor in metaphase, like the *Drosophila* orthologue Asp. A recent study showed that ASPM directly binds to the minus-ends of MTs *in vitro* using the middle region (Jiang et al., 2017), and another study on *Drosophila* Asp demonstrated that the C-terminal region is responsible for end-end interaction *in vivo* (Ito and Goshima, 2015). Thus, it is strongly suggested that cross-linking of adjacent MT minus ends is a conserved function of ASPM/Asp. However, unlike Asp in *Drosophila*, whose depletion gives rise to severe phenotypes in various cell types and causes brain size reduction as well as lethality (Ripoll et al., 1985, Saunders et al., 1997, Wakefield et al., 2001, Rujano et al., 2013, Schoborg et al., 2015), ASPM was not essential in the human HCT116 (this study) or HeLa cells (Jiang et al., 2017). Our data indicated that ASPM became indispensable for pole organisation and thereby mitotic progression when CDK5RAP2 is depleted. Normally, CDK5RAP2 and possibly other pole-focusing factor(s) are active enough to compensate for the function of ASPM in this cell type. The redundant nature of pole organisation may hold true for many other cell types in mammals. In a mouse model, mutations in *Aspm* reduce brain and testis/ovary sizes but are not lethal (Pulvers et al., 2010, Fujimori et al., 2014). A cell type-specific polar defect has been observed upon depletion of another pole-focusing factor, NuMA (Seldin et al., 2016, Silk et al., 2009). Thus, it is tempting to speculate that certain neuronal cell types in humans rely more heavily on ASPM for pole organisation than on other factors; therefore, a severe phenotype only appears in patients’ brains.

### Spindle pole focusing by CDK5RAP2

How does CDK5RAP2 depletion affect pole organisation? A possible explanation was that a reduction in centrosomal MTs affects spindle pole organisation, since they would provide a platform for pole-focusing factors such as motors. However, we could not detect gross changes in astral MT number or γ-tubulin localisation after CDK5RAP2 depletion with standard immunofluorescence microscopy, which is consistent with results from some other cell types (Chavali et al., 2016, Barr et al., 2010). Thus, although not entirely excluded, attenuation of γ-tubulin-dependent MT nucleation is less likely to account for the synthetic phenotype. A recent study showed that CDK5RAP2 binds to HSET and promotes the association of HSET with the centrosome (Chavali et al., 2016). However, HSET delocalisation is also unlikely to be the primary cause of pole disorganisation after double ASPM/CDK5RAP2 depletion in our cell line, since we could not observe a synthetic phenotype with HSET and ASPM. Moreover, HSET localisation was indistinguishable between control and CDK5RAP2-depleted cells. More recently, ASPM was shown to recruit the MT-severing protein katanin to the spindle pole in HeLa cells; the ASPM-katanin complex regulates MT minus end depolymerisation (Jiang et al., 2017). While this could also be occurring in HCT116 cells, we do not think this interaction is associated with the pole-focusing function of ASPM. Firstly, the binding site of katanin was mapped at the N-terminal region of ASPM, whereas our study using a patient mutation indicated that the loss of the C-terminal region is sufficient to produce the phenotype. Secondly, HeLa cells lacking katanin or ASPM did not exhibit a pole-unfocusing phenotype (Jiang et al., 2017). Interestingly, another recent study showed that *C. elegans* SPD-5, the functional homologue of CDK5RAP2/Cnn, forms spherical condensates *in vitro* and recruits two MAPs (XMAP215 and TPX2), and thereby tubulin, without the aid of other factors (Woodruff et al., 2017). If this were the case for CDK5RAP2, these or other MAPs on the spindle MT or MT itself might directly associate with CDK5RAP2 and support pole association of spindle MT minus ends. It would be interesting to identify the factor(s) that bridges CDK5RAP2 and MT ends.

### Spindle pole focusing and microcephaly

A model for the cause of microcephaly postulates a defect in spindle orientation and asymmetric cell division (Fish et al., 2008, Gai et al., 2016, Thornton and Woods, 2009, Lizarraga et al., 2010, Feng and Walsh, 2004, Fish et al., 2006). Consistent with this idea, spindle orientation error was reported following *ASPM* RNAi in human U2OS cells and CDK5RAP2 mutant mice (Higgins et al., 2010, Lizarraga et al., 2010). However, depletion of other proteins required for spindle orientation (LGN, aPKCλ) does not cause microcephaly in the mouse model (Megraw et al., 2011, Imai et al., 2006, Konno et al., 2008). Moreover, a recent study using a *Sas-4* mutant mouse model suggested that microcephaly is caused by mitotic delay and apoptosis, independent of spindle orientation (Insolera et al., 2014). Another recent study identified a role of ASPM in controlling centriole duplication in the mouse model, which might lead to microcephaly (Jayaraman et al., 2016). The results of the present study raise yet another possibility that spindle pole disorganisation is a cause of the severe mitotic delay that might lead to *ASPM*- and *CDK5RAP2-*linked microcephaly.

## Materials and Methods

### Plasmid construction

Plasmid information is described in Table S4. pX330-U6-Chimeric_BB-CBh-hSpCas9 (Addgene #42230 (Cong et al., 2013)) was used to construct CRISPR/Cas9 vectors according to the protocol of (Ran et al., 2013). PAM and 20-bp sgRNA sequences were selected by the Optimized CRISPR Design tool (http://crispr.mit.edu) (Table S2).

### Cell culture and line selection

The original HCT116 cell line and the HCT116 cells expressing OsTIR1 under the conditional Tet promoter and the puromycin-resistant gene (HCT116 Tet-OsTIR1) were cultured in McCoy’s 5A medium (Gibco) supplemented with 10% foetal bovine serum (Gibco) and 1% penicillin-streptomycin-amphotericin B suspension (Wako) (cells were obtained from Dr M. Kanemaki) (Natsume et al., 2016). Cells were maintained in a 37°C humid incubator with 5% CO_2_. Effectine (Qiagen) was used for plasmid transfection. For the construction of the hypomorphic *ASPM* KO line (KO^Exon1^), a CRISPR/Cas9 plasmid (pAI112) with sgRNA and SpCas9 sequences, and a donor plasmid (pAI114) harbouring the neomycin-resistant gene flanked by homologous sequences were transfected into the original HCT116 cell line. The complete *ASPM* KO line (KO^C^) was selected by co-transfection of the following two plasmids into the original HCT116 cell: a CRISPR/Cas9 plasmid (pAI119) that contains two sets of sgRNA and SpCas9 targeting the N- and C-terminal regions, and a donor plasmid (pAI118) that has the neomycin-resistant gene flanked by homologous sequences. Proper integration of the neomycin-resistant gene cassette was verified with PCR followed by sequencing. mAID-tagged cell lines expressing CDK5RAP2-mAID-mCherry were generated according to the procedures described in (Natsume et al., 2016). In brief, the HCT116 Tet-OsTIR1 line was transfected with a CRISPR/Cas9 plasmid (pTK478) and a donor plasmid (pTK472) carrying mAID-mCherry and a hygromycin-resistant gene flanked by homologous sequences. The ASPM-KO/CDK5RAP2-mAID-mCherry line was selected by co-transfection of the CRISPR/Cas9 plasmid (pAI112) and the donor plasmid (pAI114) into the CDK5RAP2-mAID-mCherry expression line. ASPM-mClover and ASPM [1-3232]-mClover lines were selected by co-transfection of the CRISPR/Cas9 plasmid (pAI113, pAI121) and the donor plasmid (pAI110, pAI120) harbouring mClover and neomycin-resistant genes flanked by homologous sequences into the CDK5RAP2-mAID-mCherry expression line. Two to three days after transfection, the cells were plated on 10-cm culture dishes and cultured in selection medium containing 200 µg/ml hygromycin (Wako), 1 µg/ml puromycin (Wako), and/or 800 µg/ml G-418 (Roche) for 10–18 days. For single colony isolation, colonies were washed with PBS, picked with a pipette tip under a microscope (EVOS XL, Thermo Fisher Scientific), and subsequently transferred to a 96-well plate containing 50 µl of Tripsin-EDTA (Natsume et al., 2016). After a few minutes, the cells were transferred to a 24-well plate containing 800 µl of the selection medium. Two to four days later, cells were transferred and grown in a 6-well plate. Half of the cells were used for genomic DNA preparation, and the rest were frozen using Bambanker or Bambanker Direct Media (Nippon Genetics). To confirm homozygous insertion, genomic DNAs were prepared from drug-resistant clones for most lines, whereas the cells were directly used as PCR templates for checking the ASPM-mClover lines (full-length and C-terminal truncation). PCR was performed using Tks Gflex DNA polymerase (TaKaRa) and corresponding primers (Table S3), as described by (Natsume et al., 2016). Proper tagging of mClover to *ASPM* was also verified by DNA sequencing after PCR.

### AID

To degrade mAID-tagged CDK5RAP2, the cells were treated with 2 µg/ml Dox (Sigma) for 24 h to induce the expression of OsTIR1, and with 500 µM IAA (Wako) for 24 h to induce the degradation of CDK5RAP2-mAID-mCherry (Natsume et al., 2016). The mCherry signal was undetectable in 78% (n = 435) of the cells treated with Dox and IAA.

### RNAi

siRNA transfection was carried out using Lipofectamine RNAiMAX (Invitrogen) and 13.3 nM siRNA. The primers for siRNA constructs are shown in Table S1 (note that we observed a possible off-target effect for at least two of the constructs). In the ASPM depletion experiment, HCT116 cells were incubated with *ASPM-*targeted or control luciferase-targeted siRNA for 42 h (live imaging) or 48 h (immunostaining). For live imaging after mAID-tagged CDK5RAP2 degradation, Dox and IAA were added to the culture at 24 h after siRNA transfection. Live imaging began at 24 h or 48 h following CDK5RAP2 or HSET siRNA treatment, respectively.

### Microscopy, image analysis, and data presentation

Microscopy was basically identical to that described in (Ito and Goshima, 2015). Live imaging was performed using spinning-disc confocal microscopy with a 60× 1.40 NA objective lens. A CSU-X1 confocal unit (Yokogawa, Japan) attached to a Nikon Ti inverted microscope, EMCCD camera ImagEM (Hamamatsu, Japan) that was cooled (−80°C) with a chiller (Julabo USA Inc.), and a stage chamber were used for image acquisition (37°C, 5% CO_2_). Characterisation of the phenotypes of fixed samples was performed with the Nikon Ti inverted microscope attached to the EMCCD camera Evolve (Roper). Three objective lenses (100× 1.49 NA, 100× lens 1.45 NA, or 40× 1.30 NA) were used. The microscopes and attached devices were controlled using Micromanager. Images were analysed with ImageJ. For live imaging, the MTs were stained with SiR-Tubulin (30 nM; Spirochrome) for >1 h prior to image acquisition. Immunostaining was performed according to the standard procedure using 6.4% paraformaldehyde, and anti-tubulin (YOL1/34, 1:500, rat monoclonal, AbD Serotec), anti-ASPM (1:1,000, rabbit polyclonal, antigen [aa 3,425–3,477], IHC-00058, Bethyl Laboratories, Inc.), and anti-HSET (1:1,000, rabbit polyclonal, A300-952A, Bethyl Laboratories, Inc.) antibodies. DNA was stained with DAPI. The fluorescence intensity of polar ASPM was measured from a single-plane image of the immunostained samples, with measurements taken from the cell’s most in-focus pole. Background intensity was subtracted from each measurement. Multiple images presented in a comparative manner in figures and movies were acquired and processed in an identical manner, except for those in Fig. 4A, in which SiR-tubulin staining in the ASPM #2 sample was stronger than the others, perhaps due to massive cell death induced by this specific siRNA; thus, the images were acquired with a shorter laser exposure for this particular sample. Graphical presentation of the data and statistical analysis were performed with the GraphPad Prism 6.0 software. The p-values were obtained after unpaired two-sided Student’s t tests. Note, however, the statistical significance of the differences as well as normality of the data distribution could not be properly discussed when sample sizes were too small (e.g. three). Nevertheless, the values served as a good indicator of the degree of change. When p-value was >0.1, we indicated ‘n.s.’ (not significant) in figures.

### Immunoblotting

Protein extracts were prepared according to a previously described protocol (Ito and Goshima, 2015). In brief, the cells were treated with the Cytobuster (EMD Millipore) solution, which also contained benzonase, dithiothreitol, and protease inhibitors (30 min on ice), followed by addition of the urea-containing solution (62.5 mM Tris, pH 6.8, 10% glycerol, 2% SDS, 4 M urea, and 2-mercaptoethanol) at room temperature (5 h). Immunoblotting was carried out with anti-CDK5RAP2 (1:10,000, rabbit polyclonal, ab70213, Abcam).

## Acknowledgements

We are grateful to Toyoaki Natsume and Masato Kanemaki (National Institute of Genetics, Japan) for reagents and technical advice, and to Momoko Nishina and Moé Yamada (Nagoya University, Japan) for technical assistance. This work was supported by Japan Society for the Promotion of Science (JSPS) KAKENHI (17H01431), the Uehara Memorial Foundation, and Takeda Science Foundation (to G.G.). A.I. was a recipient of a JSPS pre-doctoral fellowship. The authors declare no competing financial interests.

## Author contributions

Conceptualization: E.A.T., A.I., G.G.; Methodology: All authors; Investigation: E.A.T., A.I.; Writing: G.G.; Supervision: G.G.; Funding acquisition: G.G.

## Supplementary data

**Table S1.**
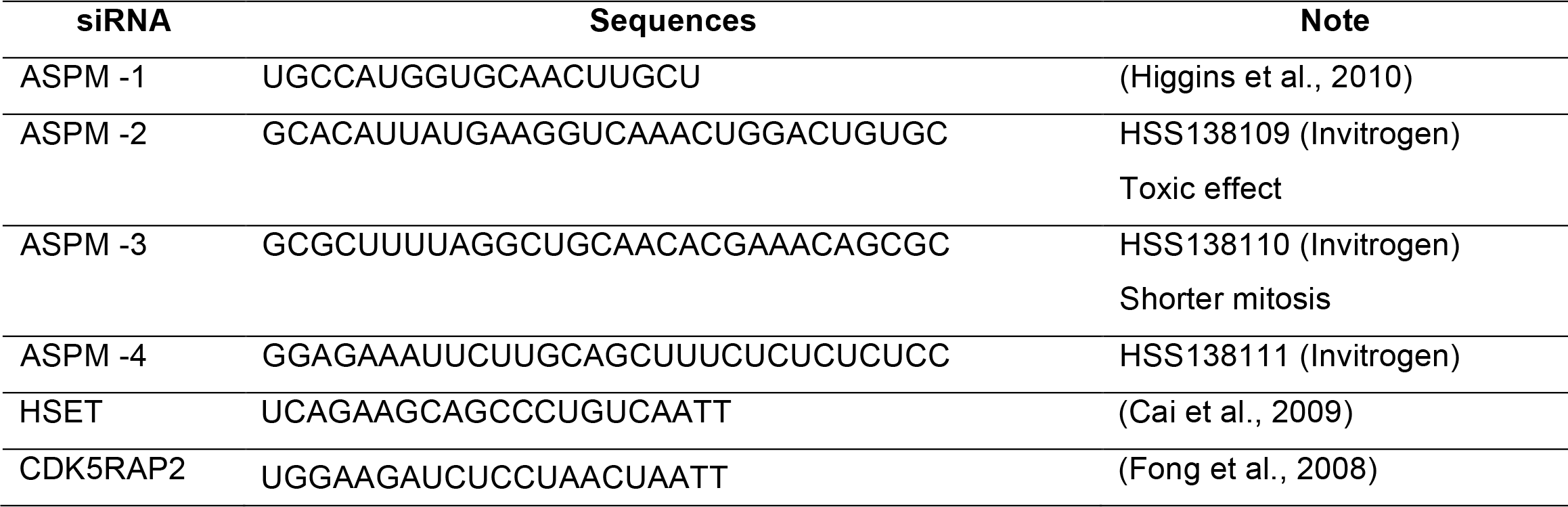
Primers for RNAi

**Table S2.**
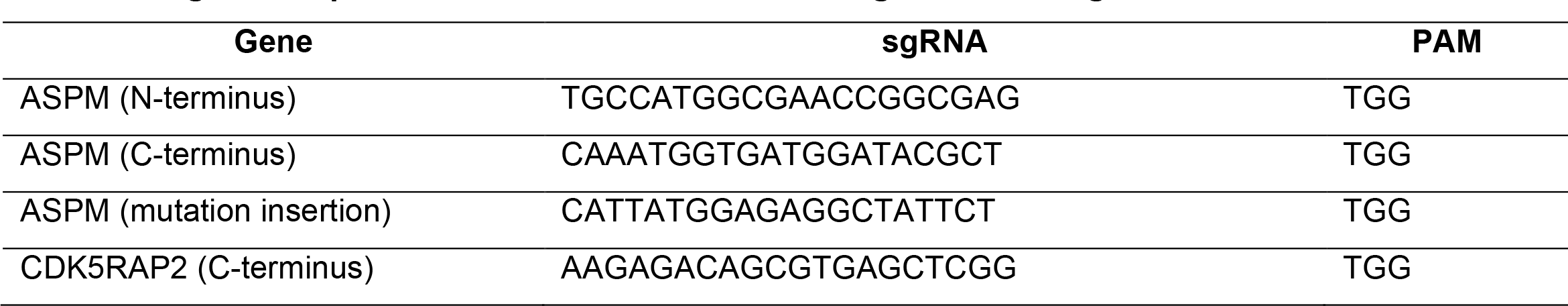
sgRNA sequences for CRISPR/Cas9-mediated genome editing

**Table S3.**
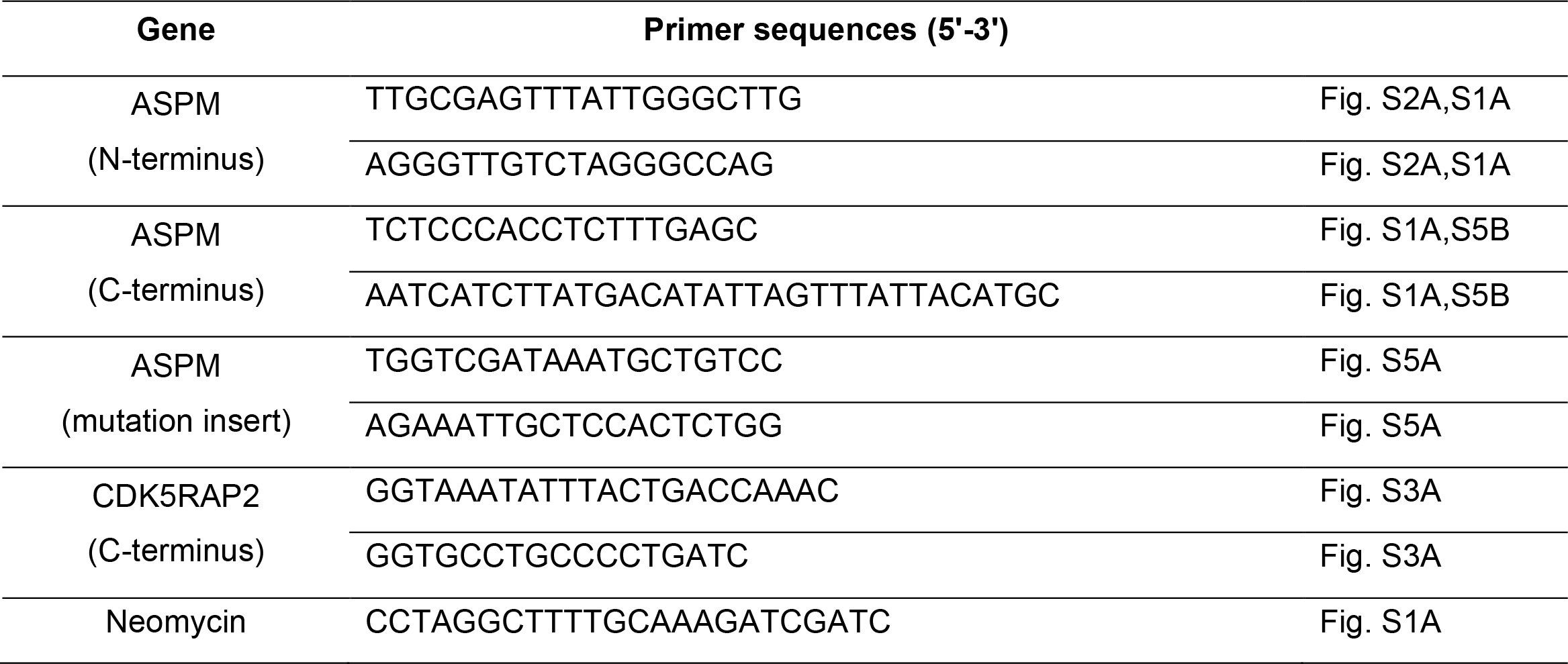
PCR primers to confirm gene editing

**Table S4.**
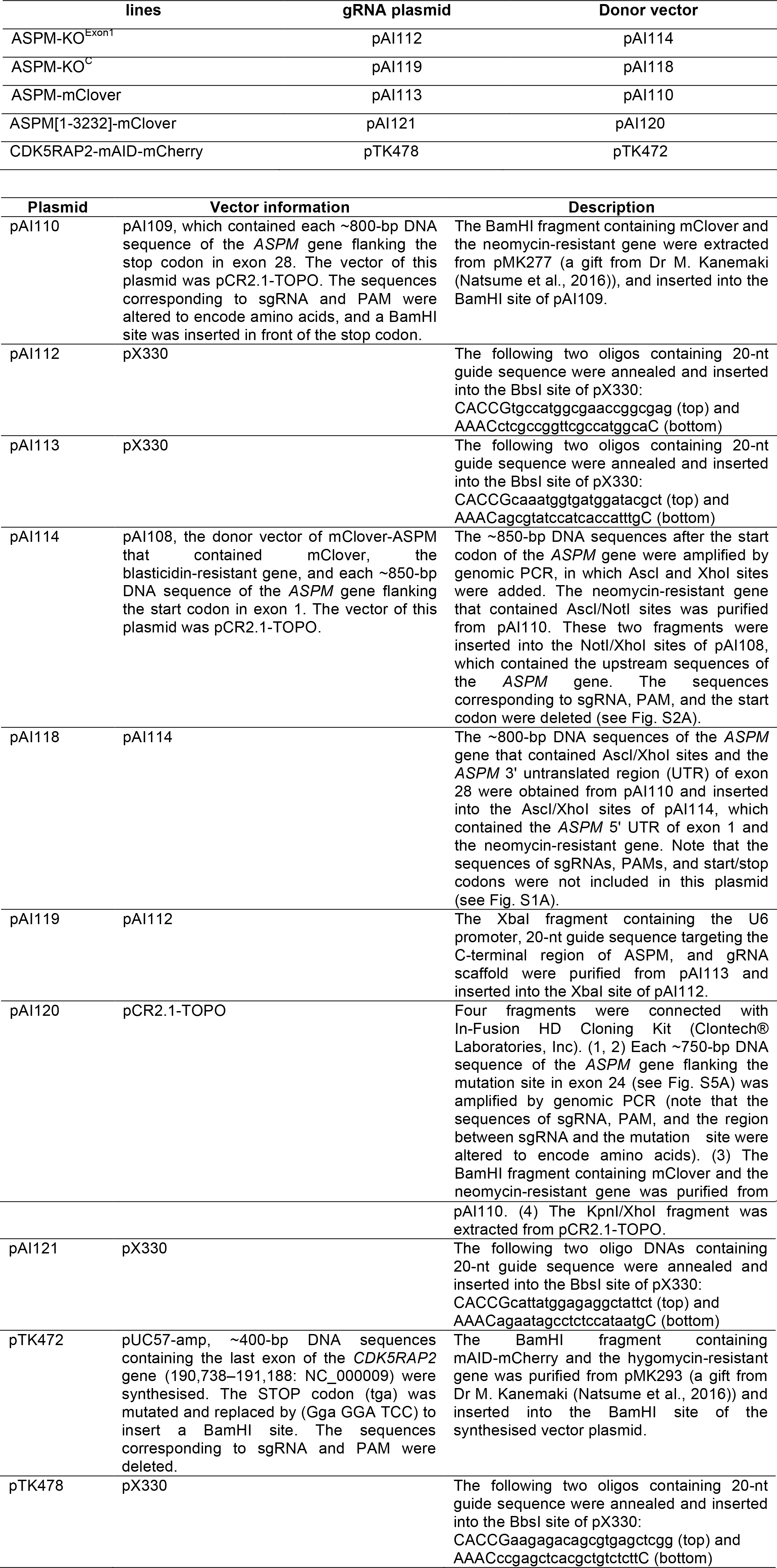
Plasmids and cell lines constructed and used in this study

**Movie 1. Normal spindle formation and mitotic progression after *ASPM* KO**

SiR-tubulin images were acquired every 3 min with spinning-disc confocal microscopy. Images were acquired with 5 z-sections (separated by 3 µm) and displayed after maximum projection. Time 0 corresponds to the timing of nuclear envelope breakdown. The images of two KO^Exon1^ lines (#4 and #23) and two complete KO^C^ lines (#25 and #27) are presented.

**Movie 2. CDK5RAP2 degradation impairs spindle pole focusing and mitotic progression in the absence of ASPM**

Images were acquired every 3 min with spinning-disc confocal microscopy. Green, CDK5RAP2-mAID-mCherry (control cell only); magenta, Sir-Tubulin. Images were acquired with 5 z-sections (separated by 3 µm) and displayed after maximum projection. Time 0 corresponds to the timing of nuclear envelope breakdown.

**Movie 3. Microcephaly mutation of ASPM impairs spindle focusing and mitotic progression in the absence of CDK5RAP2**

Images were acquired every 5 min with spinning-disc confocal microscopy. Yellow, CDK5RAP2-mAID-mCherry; magenta, Sir-Tubulin; cyan, ASPM-mClover (full-length or C-terminal truncation that mimics microcephaly mutations). Images were acquired with 5 z-sections (separated by 3 µm) and displayed after maximum projection. Time 0 corresponds to the timing of nuclear envelope breakdown.

**Figure S1.**
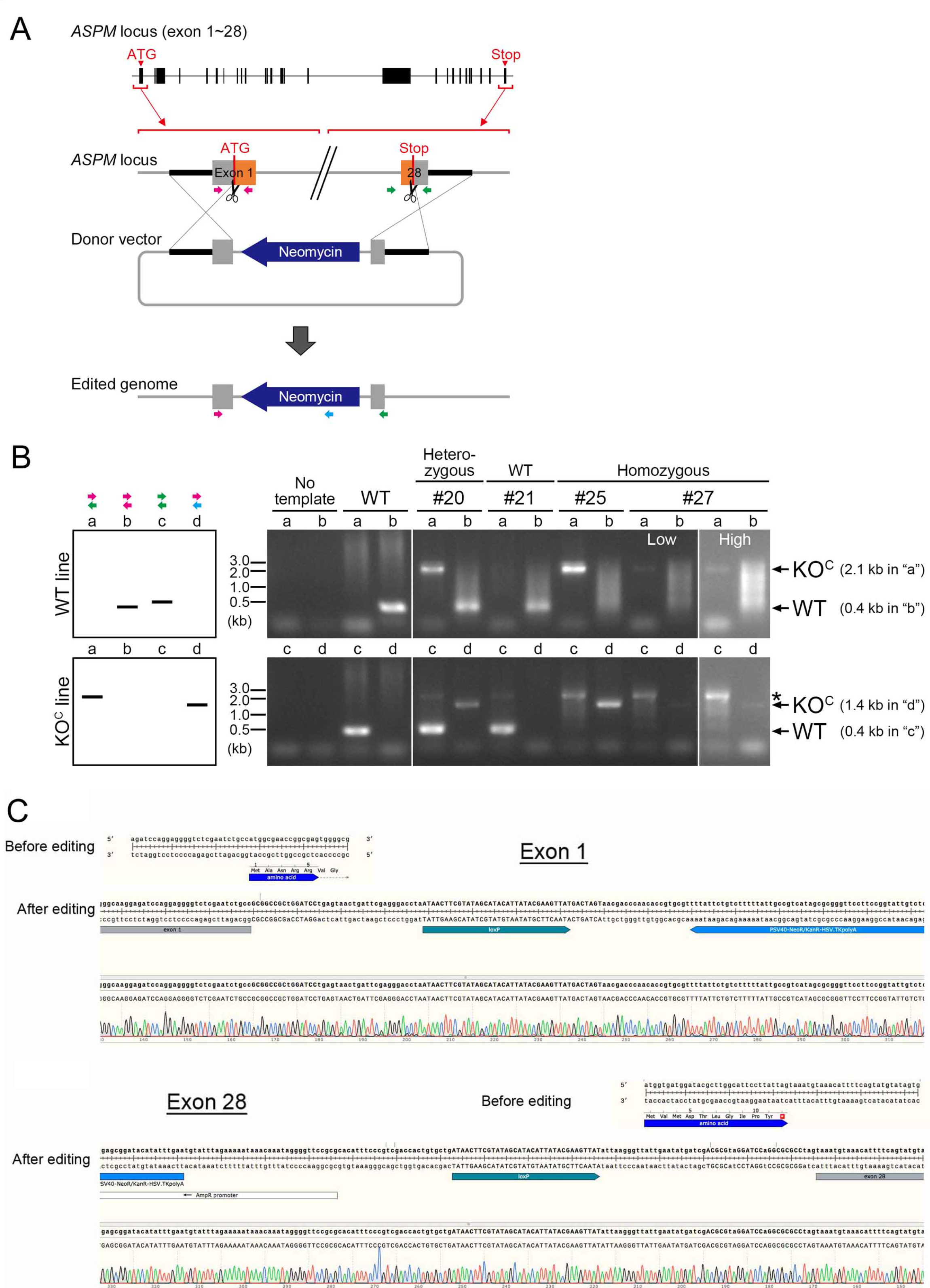
Construction of the complete *ASPM* KO line. Construction of the complete *ASPM* KO line (KOc), in which the entire *ASPM* open reading frame was replaced by the neomycin-resistant marker. Targeted integration of the marker cassette was confirmed by PCR (primers are indicated as arrows in A) and sequencing (C). Expected bands are displayed on the left of (8). Asterisk in (8) indicates a non-specific band. Note that an expected band with the primer pair ‘a’ was too long to detect.

**Figure S2.**
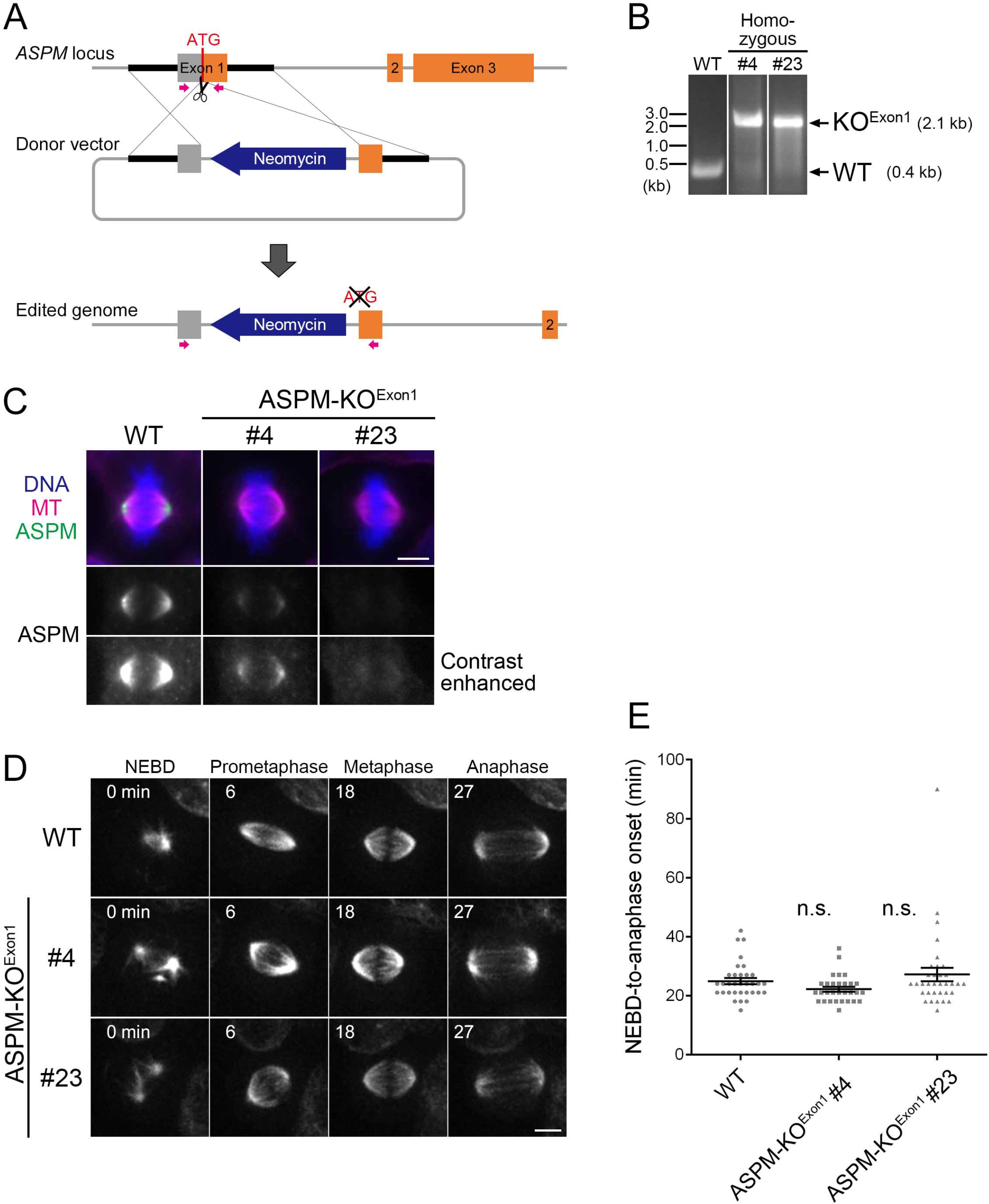
Construction of the exon 1-targeted *ASPM* KO line. **(A, B)** Construction of the hypomorphic *ASPM* KO line (KO^Exon1^ line), in which the region surrounding the start codon at the first exon of the *ASPM* gene was replaced by the neomycin-resistant marker. Targeted integration of the marker cassette was confirmed by PCR (primers are indicated as magenta arrows). **(C)** lmmunofluorescence microscopy confirmed the diminishment of the ASPM signal at the spindle pole in two selected KOExon ^1^ lines (clones #4 and #23). Note that truncated ASPM protein was still weakly expressed and detected at the spindle pole. **(D)** Spindle dynamics of *ASPM* KOExon ^1^ li nes. MTs were visualized by SiR-tubulin staining. Images were acquired with 5 z-sections (separated by 3 µm) and displayed after maximum projection. No abnormality was detected. See also Movie 1. **(E)** Mitotic duration with or without ASPM. Data represent the mean± SEM. WT; 25 ± 1 (n 33), KO # 4; 22 ± 1 (n 30, p > 0.05), KO #23; 27 ± 2 (n = 34, p > 0.3). n.s. stands for ‘not significant’. Bars, 5 µm.

**Figure S3.**
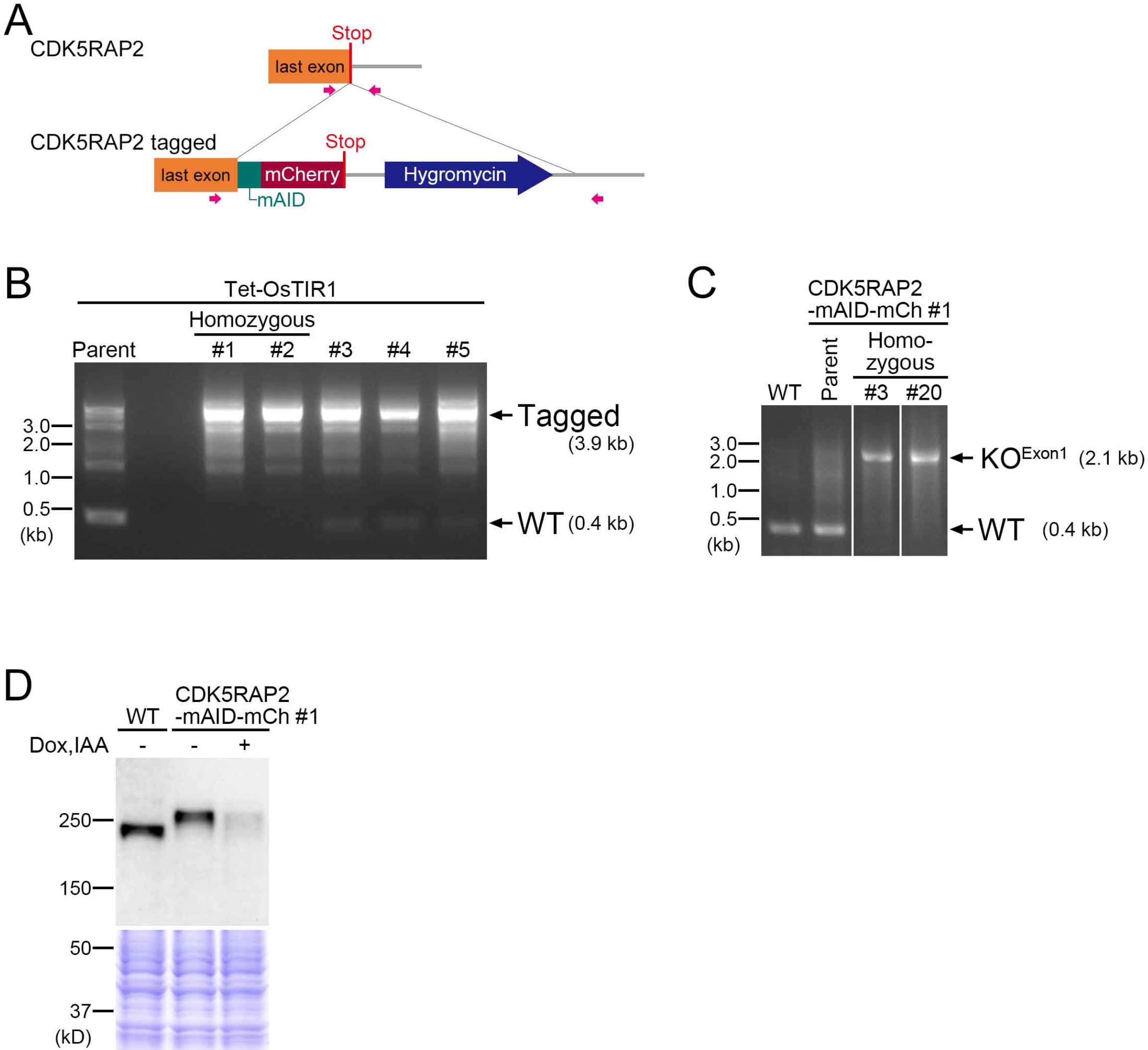
Construction of the CDK5RAP2-mAID-mCherry (degron) line. **(A, B)** Targeted integration of the marker cassette was confirmed by PCR (primers are indicated as magenta arrows). Clone #1 was used as the parent line of the following experiments. **(C)** ASPM was knocked out with the strategy described in Fig. S2A. **(D)** Confirmation of CDK5RAP2-mAID-mCherry protein degradation by Dox and IAA treatment. lmmunoblotting using anti-CDK5RAP2 antibody and Coomassie staining of whole-cell extracts are presented. This blotting also confirmed the homozygous tagging of mAID-mCherry to CDK5RAP2.

**Figure S4.**
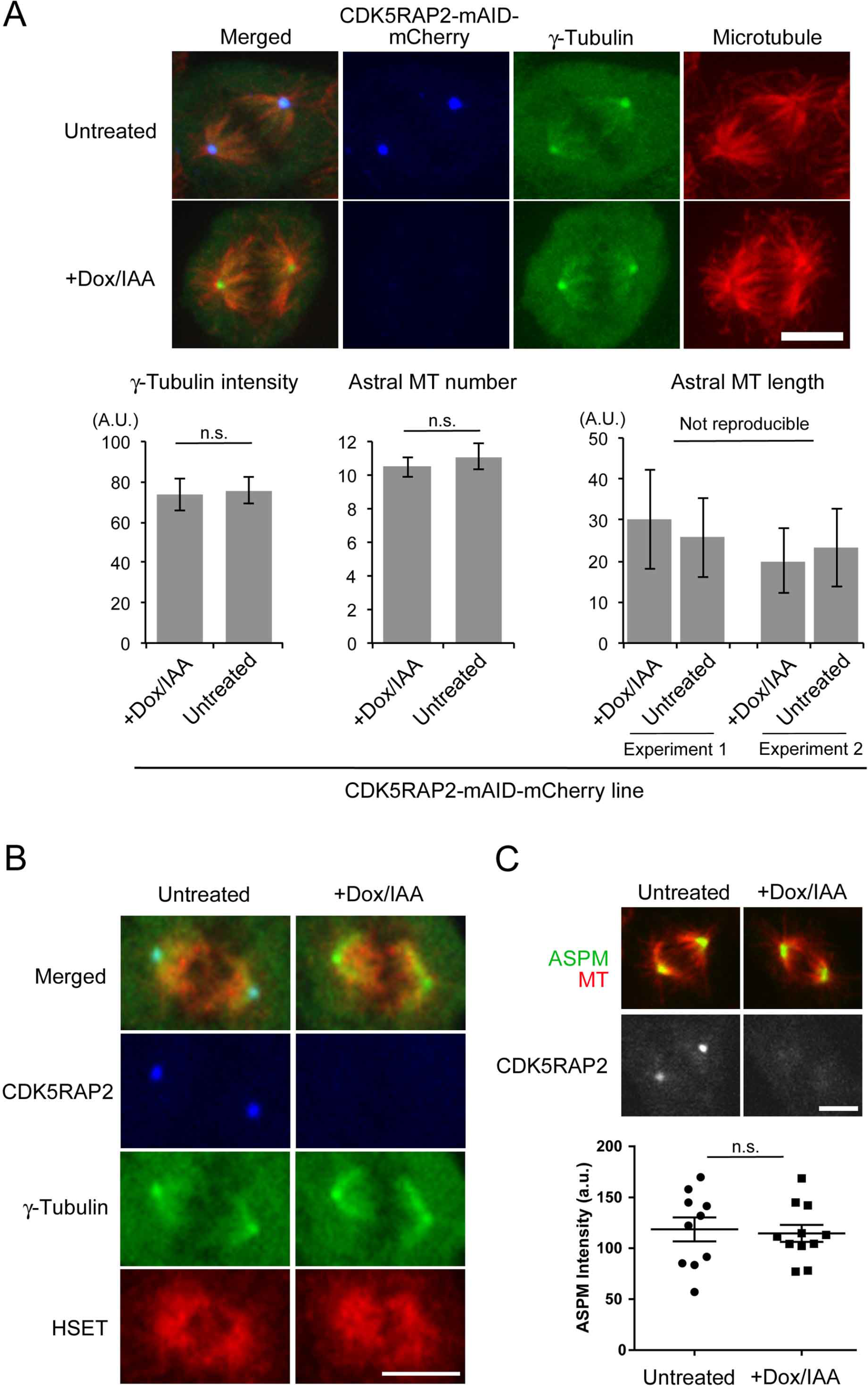
Effect of CDK5RAP2 depletion on astral MTs, y-tubulin, HSET, and ASPM localisation. (A) Quantification of y-tubulinsignal intensity (±SEM, 10 cells each), MT number (±SEM, 20 asters each), and MT length (±SD, n = 117, 126, 93, 96 from left to right) after immunostaining in the presence (untreated) or absence (+Dox/lAA) of CDK5RAP2. Astral MT numbers and length were counted based on the projected image, whereas a single focal plane with the most in-focus centrosome was selected for y-tubulinintensity measurement. n.s. stands for ‘not significant’ (p > 0.5). **(B)** Mitotic localisation of HSET revealed by anti-HSET immunostaining was indistinguishable between control and CDK5RAP2-depleted cells. **(C)** A similar level of pole accumulation of ASPM (anti-ASPM staining) in the absence of CDK5RAP2 (p > 0.78, n = 10 each). Bars, 5 µm.

**Figure S5.**
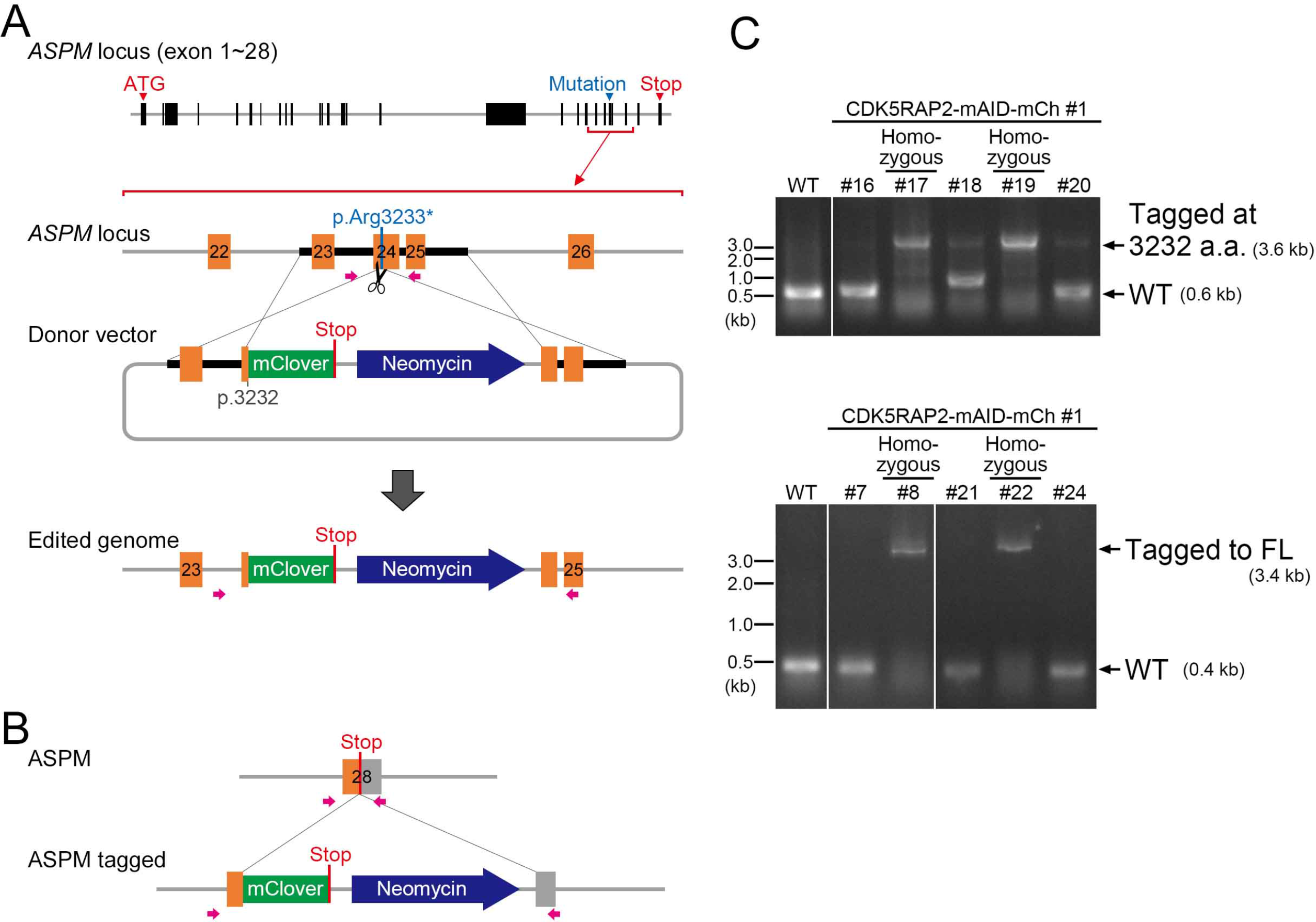
Construction of full-length and truncated ASPM-mClover lines. (A) Construction of the truncated ASPM line, in which mClover and the neomycin-resistant gene were inserted at exon 24, which mimicked the second-most C-terminally truncated protein identified among microcephaly patients (aa 1-3,232). **(B)** As a control, mClover and the marker were integrated at the C-terminus of the full-length ASPM protein. **(C)** Targeted integration of mClover and the marker cassette was confirmed by PCR (primers are indicated as magenta arrows in A and B). The CDK5RAP2-mAID-mCherry line (clone #1; Fig. S3B) was used as the host of integration. Proper tagging of mClover (ASPM-mClover #8 and #22, ASPM [1-3,232 a.a.]-mClover#17 and #19) was also verified by DNA sequencing after PCR.

## References

Abdel-Hamid, M. S., Ismail, M. F., Darwish, H. A., Effat, L. K., Zaki, M. S. and Abdel-Salam, G. M. (2016). Molecular and phenotypic spectrum of ASPM-related primary microcephaly: Identification of eight novel mutations. Am J Med Genet A. 170, 2133–2140.

Barr, A. R., Kilmartin, J. V. and Gergely, F. (2010). CDK5RAP2 functions in centrosome to spindle pole attachment and DNA damage response. J Cell Biol. 189, 23–39.

Baumbach, J., Novak, Z. A., Raff, J. W. and Wainman, A. (2015). Dissecting the function and assembly of acentriolar microtubule organizing centers in Drosophila cells in vivo. PLoS Genet. 11, e1005261.

Bond, J., Roberts, E., Mochida, G. H., Hampshire, D. J., Scott, S., Askham, J. M., Springell, K., Mahadevan, M., Crow, Y. J., Markham, A. F., Walsh, C. A. and Woods, C. G. (2002). ASPM is a major determinant of cerebral cortical size. Nat Genet. 32, 316–320.

Bond, J., Roberts, E., Springell, K., Lizarraga, S. B., Scott, S., Higgins, J., Hampshire, D. J., Morrison, E. E., Leal, G. F., Silva, E. O., Costa, S. M., Baralle, D., Raponi, M., Karbani, G., Rashid, Y., Jafri, H., Bennett, C., Corry, P., Walsh, C. A. and Woods, C. G. (2005). A centrosomal mechanism involving CDK5RAP2 and CENPJ controls brain size. Nat Genet. 37, 353–355.

Bond, J., Scott, S., Hampshire, D. J., Springell, K., Corry, P., Abramowicz, M. J., Mochida, G. H., Hennekam, R. C., Maher, E. R., Fryns, J. P., Alswaid, A., Jafri, H., Rashid, Y., Mubaidin, A., Walsh, C. A., Roberts, E. and Woods, C. G. (2003). Protein-truncating mutations in ASPM cause variable reduction in brain size. Am J Hum Genet. 73, 1170–1177.

Cai, S., O’connell, C. B., Khodjakov, A. and Walczak, C. E. (2009). Chromosome congression in the absence of kinetochore fibres. Nat Cell Biol. 11, 832–838.

Chavali, P. L., Chandrasekaran, G., Barr, A. R., Tatrai, P., Taylor, C., Papachristou, E. K., Woods, C. G., Chavali, S. and Gergely, F. (2016). A CEP215-HSET complex links centrosomes with spindle poles and drives centrosome clustering in cancer. Nat Commun. 7, 11005.

Choi, Y. K., Liu, P., Sze, S. K., Dai, C. and Qi, R. Z. (2010). CDK5RAP2 stimulates microtubule nucleation by the gamma-tubulin ring complex. J Cell Biol. 191, 1089–1095.

Cong, L., Ran, F. A., Cox, D., Lin, S., Barretto, R., Habib, N., Hsu, P. D., Wu, X., Jiang, W., Marraffini, L. A. and Zhang, F. (2013). Multiplex genome engineering using CRISPR/Cas systems. Science. 339, 819–823.

Feng, Y. and Walsh, C. A. (2004). Mitotic spindle regulation by Nde1 controls cerebral cortical size. Neuron. 44, 279–293.

Fish, J. L., Dehay, C., Kennedy, H. and Huttner, W. B. (2008). Making bigger brains-the evolution of neural-progenitor-cell division. J Cell Sci. 121, 2783–2793.

Fish, J. L., Kosodo, Y., Enard, W., Paabo, S. and Huttner, W. B. (2006). Aspm specifically maintains symmetric proliferative divisions of neuroepithelial cells. Proc Natl Acad Sci U S A. 103, 10438–10443.

Fong, K. W., Choi, Y. K., Rattner, J. B. and Qi, R. Z. (2008). CDK5RAP2 is a pericentriolar protein that functions in centrosomal attachment of the gamma-tubulin ring complex. Mol Biol Cell. 19, 115–125.

Fujimori, A., Itoh, K., Goto, S., Hirakawa, H., Wang, B., Kokubo, T., Kito, S., Tsukamoto, S. and Fushiki, S. (2014). Disruption of Aspm causes microcephaly with abnormal neuronal differentiation. Brain Dev. 36, 661–669.

Gai, M., Bianchi, F. T., Vagnoni, C., Verni, F., Bonaccorsi, S., Pasquero, S., Berto, G. E., Sgro, F., Chiotto, A. M., Annaratone, L., Sapino, A., Bergo, A., Landsberger, N., Bond, J., Huttner, W. B. and Di Cunto, F. (2016). ASPM and CITK regulate spindle orientation by affecting the dynamics of astral microtubules. EMBO Rep. 17, 1396–1409.

Goshima, G., Nedelec, F. and Vale, R. D. (2005). Mechanisms for focusing mitotic spindle poles by minus end-directed motor proteins. J Cell Biol. 171, 229–240.

Heald, R., Tournebize, R., Habermann, A., Karsenti, E. and Hyman, A. (1997). Spindle assembly in Xenopus egg extracts: respective roles of centrosomes and microtubule self-organization. J Cell Biol. 138, 615–628.

Higgins, J., Midgley, C., Bergh, A. M., Bell, S. M., Askham, J. M., Roberts, E., Binns, R. K., Sharif, S. M., Bennett, C., Glover, D. M., Woods, C. G., Morrison, E. E. and Bond, J. (2010). Human ASPM participates in spindle organisation, spindle orientation and cytokinesis. BMC Cell Biol. 11, 85.

Imai, F., Hirai, S., Akimoto, K., Koyama, H., Miyata, T., Ogawa, M., Noguchi, S., Sasaoka, T., Noda, T. and Ohno, S. (2006). Inactivation of aPKClambda results in the loss of adherens junctions in neuroepithelial cells without affecting neurogenesis in mouse neocortex. Development. 133, 1735–1744.

Insolera, R., Bazzi, H., Shao, W., Anderson, K. V. and Shi, S. H. (2014). Cortical neurogenesis in the absence of centrioles. Nat Neurosci. 17, 1528–1535.

Ito, A. and Goshima, G. (2015). Microcephaly protein Asp focuses the minus ends of spindle microtubules at the pole and within the spindle. J Cell Biol. 211, 999–1009.

Jayaraman, D., Kodani, A., Gonzalez, D. M., Mancias, J. D., Mochida, G. H., Vagnoni, C., Johnson, J., Krogan, N., Harper, J. W., Reiter, J. F., Yu, T. W., Bae, B. I. and Walsh, C. A. (2016). Microcephaly Proteins Wdr62 and Aspm Define a Mother Centriole Complex Regulating Centriole Biogenesis, Apical Complex, and Cell Fate. Neuron. 92, 813–828.

Jiang, K., Rezabkova, L., Hua, S., Liu, Q., Capitani, G., Altelaar, A. F., Heck, A. J. R., Kammerer, R. A., Steinmetz, M. O. and Akhmanova, A. (2017). Microtubule minus-end regulation at spindle poles by an ASPM-katanin complex. Nat Cell Biol. 19, 480–492.

Konno, D., Shioi, G., Shitamukai, A., Mori, A., Kiyonari, H., Miyata, T. and Matsuzaki, F. (2008). Neuroepithelial progenitors undergo LGN-dependent planar divisions to maintain self-renewability during mammalian neurogenesis. Nat Cell Biol. 10, 93–101.

Kouprina, N., Pavlicek, A., Collins, N. K., Nakano, M., Noskov, V. N., Ohzeki, J., Mochida, G. H., Risinger, J. I., Goldsmith, P., Gunsior, M., Solomon, G., Gersch, W., Kim, J. H., Barrett, J. C., Walsh, C. A., Jurka, J., Masumoto, H. and Larionov, V. (2005). The microcephaly ASPM gene is expressed in proliferating tissues and encodes for a mitotic spindle protein. Hum Mol Genet. 14, 2155–2165.

Lizarraga, S. B., Margossian, S. P., Harris, M. H., Campagna, D. R., Han, A. P., Blevins, S., Mudbhary, R., Barker, J. E., Walsh, C. A. and Fleming, M. D. (2010). Cdk5rap2 regulates centrosome function and chromosome segregation in neuronal progenitors. Development. 137, 1907–1917.

Megraw, T. L., Kao, L. R. and Kaufman, T. C. (2001). Zygotic development without functional mitotic centrosomes. Curr Biol. 11, 116–120.

Megraw, T. L., Sharkey, J. T. and Nowakowski, R. S. (2011). Cdk5rap2 exposes the centrosomal root of microcephaly syndromes. Trends Cell Biol. 21, 470–480.

Morales-Mulia, S. and Scholey, J. M. (2005). Spindle pole organization in Drosophila S2 cells by dynein, abnormal spindle protein (Asp), and KLP10A. Mol Biol Cell. 16, 3176–3186.

Mountain, V., Simerly, C., Howard, L., Ando, A., Schatten, G. and Compton, D. A. (1999). The kinesin-related protein, HSET, opposes the activity of Eg5 and cross-links microtubules in the mammalian mitotic spindle. J Cell Biol. 147, 351–366.

Natsume, T., Kiyomitsu, T., Saga, Y. and Kanemaki, M. T. (2016). Rapid Protein Depletion in Human Cells by Auxin-Inducible Degron Tagging with Short Homology Donors. Cell Rep. 15, 210–218.

Nicholas, A. K., Swanson, E. A., Cox, J. J., Karbani, G., Malik, S., Springell, K., Hampshire, D., Ahmed, M., Bond, J., Di Benedetto, D., Fichera, M., Romano, C., Dobyns, W. B. and Woods, C. G. (2009). The molecular landscape of ASPM mutations in primary microcephaly. J Med Genet. 46, 249–253.

Nishimura, K., Fukagawa, T., Takisawa, H., Kakimoto, T. and Kanemaki, M. (2009). An auxin-based degron system for the rapid depletion of proteins in nonplant cells. Nat Methods. 6, 917–922.

Paramasivam, M., Chang, Y. J. and Loturco, J. J. (2007). ASPM and citron kinase co-localize to the midbody ring during cytokinesis. Cell Cycle. 6, 1605–1612.

Pulvers, J. N., Bryk, J., Fish, J. L., Wilsch-Brauninger, M., Arai, Y., Schreier, D., Naumann, R., Helppi, J., Habermann, B., Vogt, J., Nitsch, R., Toth, A., Enard, W., Paabo, S. and Huttner, W. B. (2010). Mutations in mouse Aspm (abnormal spindle-like microcephaly associated) cause not only microcephaly but also major defects in the germline. Proc Natl Acad Sci U S A. 107, 16595–16600.

Ran, F. A., Hsu, P. D., Wright, J., Agarwala, V., Scott, D. A. and Zhang, F. (2013). Genome engineering using the CRISPR-Cas9 system. Nat Protoc. 8, 2281–2308.

Ripoll, P., Pimpinelli, S., Valdivia, M. M. and Avila, J. (1985). A cell division mutant of Drosophila with a functionally abnormal spindle. Cell. 41, 907–912.

Rujano, M. A., Sanchez-Pulido, L., Pennetier, C., Le Dez, G. and Basto, R. (2013). The microcephaly protein Asp regulates neuroepithelium morphogenesis by controlling the spatial distribution of myosin II. Nat Cell Biol. 15, 1294–1306.

Saunders, R. D., Avides, M. C., Howard, T., Gonzalez, C. and Glover, D. M. (1997). The Drosophila gene abnormal spindle encodes a novel microtubule-associated protein that associates with the polar regions of the mitotic spindle. J Cell Biol. 137, 881–890.

Schoborg, T., Zajac, A. L., Fagerstrom, C. J., Guillen, R. X. and Rusan, N. M. (2015). An Asp-CaM complex is required for centrosome-pole cohesion and centrosome inheritance in neural stem cells. J Cell Biol. 211, 987–998.

Seldin, L., Muroyama, A. and Lechler, T. (2016). NuMA-microtubule interactions are critical for spindle orientation and the morphogenesis of diverse epidermal structures. Elife. 5.

Silk, A. D., Holland, A. J. and Cleveland, D. W. (2009). Requirements for NuMA in maintenance and establishment of mammalian spindle poles. J Cell Biol. 184, 677–690.

Tan, C. A., Del Gaudio, D., Dempsey, M. A., Arndt, K., Botes, S., Reeder, A. and Das, S. (2014). Analysis of ASPM in an ethnically diverse cohort of 400 patient samples: perspectives of the molecular diagnostic laboratory. Clin Genet. 85, 353–358.

Thornton, G. K. and Woods, C. G. (2009). Primary microcephaly: do all roads lead to Rome? Trends Genet. 25, 501–510.

Wakefield, J. G., Bonaccorsi, S. and Gatti, M. (2001). The drosophila protein asp is involved in microtubule organization during spindle formation and cytokinesis. J Cell Biol. 153, 637–648.

Wong, Y. L., Anzola, J. V., Davis, R. L., Yoon, M., Motamedi, A., Kroll, A., Seo, C. P., Hsia, J. E., Kim, S. K., Mitchell, J. W., Mitchell, B. J., Desai, A., Gahman, T. C., Shiau, A. K. and Oegema, K. (2015). Cell biology. Reversible centriole depletion with an inhibitor of Polo-like kinase 4. Science. 348, 1155–1160.

Woodruff, J. B., Ferreira Gomes, B., Widlund, P. O., Mahamid, J., Honigmann, A. and Hyman, A. A. (2017). The Centrosome Is a Selective Condensate that Nucleates Microtubules by Concentrating Tubulin. Cell. 169, 1066–1077 e1010.

Zhong, X., Liu, L., Zhao, A., Pfeifer, G. P. and Xu, X. (2005). The abnormal spindle-like, microcephaly-associated (ASPM) gene encodes a centrosomal protein. Cell Cycle. 4, 1227–1229.

